# Sex differences in seizure presentation in Dravet syndrome model mice, *Scn1a^+/-^*

**DOI:** 10.1101/2023.08.10.552872

**Authors:** Sheryl Anne D. Vermudez, Rui Lin, Gabrielle McGinty, Amanda Liebhardt, Yongho Choe, Benjamin Hui, Ella Lubbers, Sameer C. Dhamne, Mustafa Q. Hameed, Alexander Rotenberg

**Author notes:** **Corresponding author** Alexander Rotenberg, 300 Longwood Avenue, Fegan 9, BCH 3063, Boston, MA 02115.

## Abstract

Dravet syndrome (DS) is an epileptic encephalopathy mostly due to haploinsufficiency of the *SCN1A* voltagegated sodium channel subunit. Disease presentation (i.e., severe seizures and early life mortality) is highly recapitulated in mice haploinsufficient in *Scn1a* (*Scn1a*^*+/-*^). However, phenotypic characterization in *Scn1a*^*+/-*^ mice in a sex and temporal manner is limited. Given the reliance of mouse models for studying disease pathophysiology and for the development of novel treatments, we tested whether mortality and seizure morbidity differed among young and adult male and female *Scn1a*^*+/-*^ animals in the F1 hybrid C57×129S6 background. We found increased mortality in female *Scn1a*^*+/-*^ mice regardless of age compared to their male counterparts (n = 120-125 mice/sex; *p < 0*.*05*). Interestingly, long-term video EEG recordings revealed the opposite for morbidity as seizure frequency and severity were escalated in adult male *Scn1a*^*+/-*^ animals (n = 21-30 mice/sex; *p < 0*.*05* or *p < 0*.*01*). Adult female *Scn1a*^*+/-*^ mice, however, are more hyperactive (*p < 0*.*05*), which could be related to sleep impairment and contribute to the increased mortality despite decreased seizure morbidity. Overall, the phenotypic presentation of *Scn1a*^*+/-*^ mice is sex-dependent and may have translational implications for therapeutic drug discovery and basic biology understanding in DS.

**Short Summary:** Sex differences in mortality and seizure morbidity are discovered in *Scn1a* haploinsufficient mice, *Scn1a*^*+/-*^, which faithfully model the epileptic encephalopathy disorder, Dravet syndrome (DS). Female *Scn1a*^*+/-*^ mice die more across all ages, whereas adult male *Scn1a*^*+/-*^ mice have more seizures that are of greater severity. Hyperactivity, as a proxy for sleep disruption, may contribute to the increased mortality in female *Scn1a*^*+/-*^ mice despite decreased seizure incidence and severity. These sex-specific findings may have considerable impact in therapeutic discovery and development for DS and other *SCN1A*-related disorders.

## Introduction

Haploinsufficiency in *SCN1A*, a gene encoding the voltage-gated sodium channel Nav1.1, accounts for about 80% of Dravet syndrome (DS) (1,2). DS is a monogenic epilepsy that commonly first presents as febrile seizures in young infants and is refractory to most anti-epileptic drugs. Interestingly, clinical observations indicate that DS incidence and mortality may be sex-dependent. Several reports show that DS incidence is more prominent in females with female:male ratios of 1.11:1 – 1:27:1 (1–4). Deaths related to sudden unexpected death in epilepsy (SUDEP) or status epilepticus are also higher in female patients with DS (3,5).

*Scn1a* haploinsufficient mutant mice, such as the *Scn1a(+/-)*^*tm1Kea*^ model (6–8), recapitulates the major DS symptoms of epilepsy and early life mortality. SUDEP incidence in DS patients is prevalent during the early phases of the disease, known as febrile and worsening stages; however, despite the decreased frequency of seizures, SUDEP remains high during the next phase called stabilization stage (9). The stabilization stage is characterized by reduced seizure frequency and severity. Only a handful of studies with *Scn1a* mutant mice focus during this stabilization stage, and even so, only in young adult mice, where seizures and SUDEP are less frequent (10). We recently reported that adult *Scn1a*^*+/-*^ mice also exhibit spontaneous seizures (11), albeit again not as frequent as in juvenile mice. Yet, the influence of age and sex on SUDEP and seizure characteristics have not been fully described in these rodent models. Recent studies with juvenile *Scn1a* mutant mice demonstrate that female *Scn1a* mutant mice die earlier than their male counterparts (10,12). These findings have implications for assessing treatment efficacy as shown previously with female mice responding better to gene therapy (12). Therefore, there is a need to uncover sex variations across age in the *Scn1a* mutant mice.

Here, we aimed to characterize the sex differences of the epilepsy phenotype in adult DS model mice, *Scn1a(+/-)*^*tm1Kea*^ on the F1 hybrid C57×129S6 background, hereby referred to as *Scn1a*^*+/-*^ mice. Our results demonstrate premature mortality is biased to female *Scn1a*^*+/-*^ mice both at young (as previously shown) and adult (a novel finding) ages. A switch in sex bias occurs with morbidity as seizure frequency and severity are increased more in male *Scn1a*^*+/-*^ mice. At this adult age, female *Scn1a*^*+/-*^ mice are more hyperactive, suggesting increased wakefulness. The persistent higher mortality risk in female *Scn1a*^*+/-*^ mice despite a decline in seizure morbidity thus may in part be due to sleep impairment. Overall, these novel findings of sex-dependent disease presentation in adult *Scn1a*^*+/-*^ mice may have implications in the understanding of DS and *SCN1A* pathophysiology, as well as in novel therapeutic discovery.

## Materials & Methods

### Animals

All experiments were approved by the Institutional Animal Care and Use Committee at Boston Children’s Hospital and in accordance with the NIH Guide for the Care and Use of Laboratory Animals and the United States Public Health Service’s Policy on Humane Care and Use of Laboratory Animals. 129S-*Scn1a*^*tm1Kea*^/Mmjax males (MMRRC stock #037107-JAX) (8) were bred to wildtype (WT) 129S1/SvImJ females (JAX stock #002448) to maintain a live colony of 129S *Scn1a*^*+/-*^ heterozygotes. 129S HET males were then crossed with WT C57BL/6J female mice (JAX stock #000664) to generate F1 hybrid (C57×129S) (*Scn1a*^*+/+*^) and *Scn1a*^*+/-*^ heterozygous animals. All experiments were performed in mixed-sex, age-matched cohorts.

### Video Electroencephalography (video-EEG)

Adult P83 mice underwent wireless telemetry transmitter (ETA-F10, DSI, MN) implantation with skull screw electrodes (active: right parietal cortex (relative to Bregma: AP-0.42cm; ML-0.28cm); reference: left olfactory bulb (relative to Bregma: AP+0.15cm; ML+0.15cm)) as described previously (11). After 1 week of post-operative recovery (at P90), 1 week of epidural 1-channel continuous video-EEG (sampling rate: 1 kHz) and home-cage locomotion activity (sampling rate: 0.1 Hz) were recorded from each mouse (Ponemah Software v6.51, DSI, MN).

### Data Analyses

Mortality was calculated across multiple litters through age P90. Video-EEG data were scored for seizures using a semi-automated seizure detection algorithm (NeuroScore 3.4.1, DSI, MN) where automatically marked events were verified by visual review of real-time video and spectrogram (11). Generalized tonic-clonic (GTC) seizure frequency, total seizure time (ictal time), and mean seizure duration were calculated. Seizures and activity counts were categorized into 3 time periods – full 24 hours, 12-hour light (7 am – 7 pm), and 12-hour dark (7 pm – 7am) – to study circadian fluctuations in seizure and activity presentation.

### Statistical Analyses

Statistics were carried out using Prism 9 (GraphPad) and Excel (Microsoft). All data shown represent mean ± SEM. Statistical significance between groups was determined using Log-rank test, Chi-square test, simple linear regression, or unpaired t-tests. Sample size (denoted as “n”), statistical test and results of statistical analyses are specified in each figure legend.

## Results

Early mortality has been previously observed in *Scn1a* haploinsufficient mouse models (10,12); however, sex differences have not been fully characterized especially with respect to age. In addition to reproducing previous data of *Scn1a*^*+/-*^ mice displaying increased mortality than WT littermates (6,8), we observed excess female *Scn1a*^*+/-*^ mouse mortality (Fig 1A-B). Increased female mortality was prominent both in juvenile (<P30), as previously observed (10,12), and adult stages (P30-90). Moreover, linear regression analyses identified that female *Scn1a*^*+/-*^ mice display enhanced mortality throughout life, specifically at ages P18-30, P30-60, P60-90 and P80-90 compared to male *Scn1a*^*+/-*^ mice (Fig 1C).

**Figure 1:**
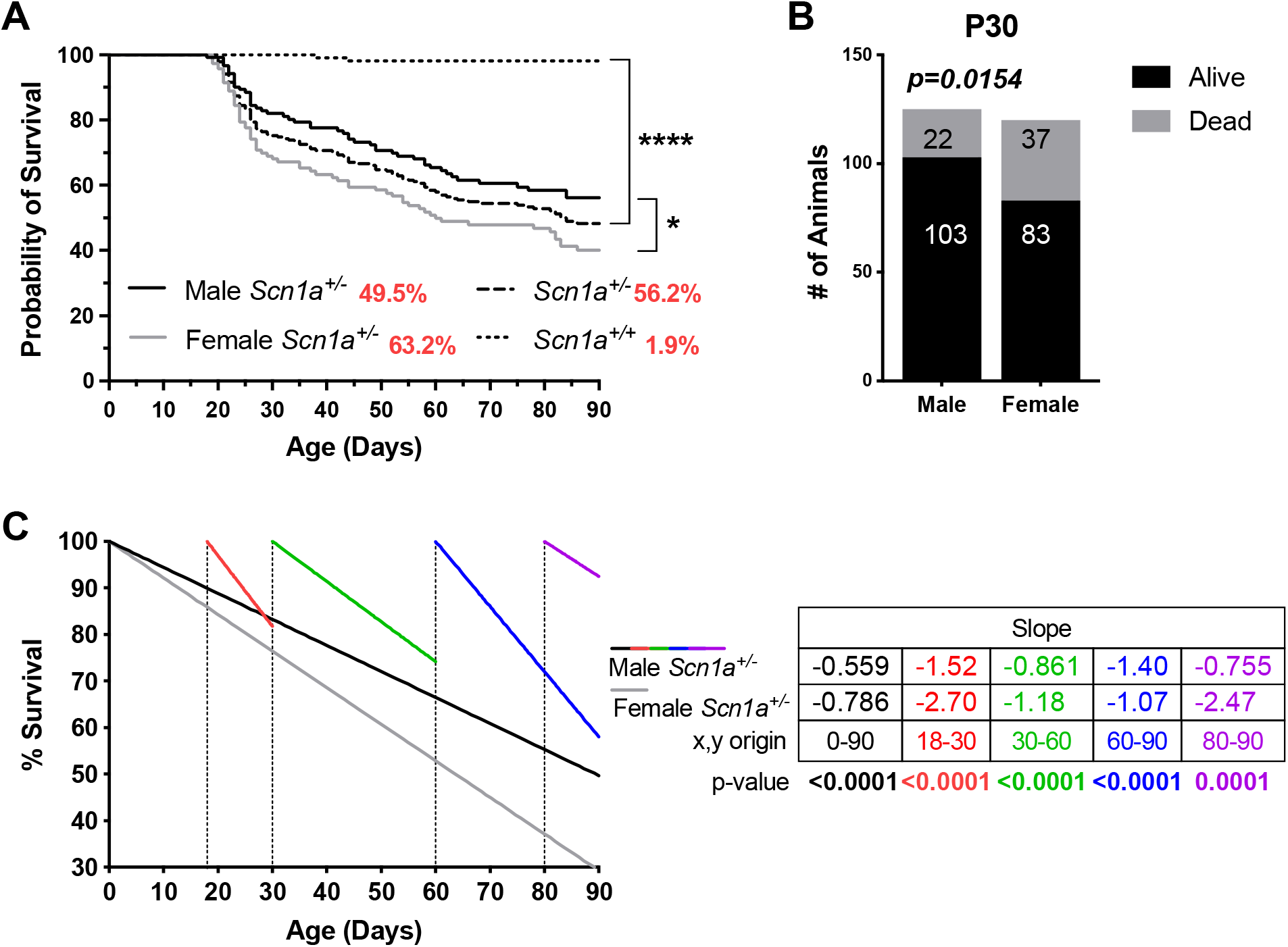
Increased mortality in female *Scn1a*^*+/-*^ mice. **A)** Kaplan-Meier curve displaying increased mortality in *Scn1a*^*+/-*^ mice compared to WT littermates (*Scn1a*^*+/+*^), where no mortality is observed. Mortality is more prevalent in female *Scn1a*^*+/-*^ mice than their male counterparts. % mortality indicated in red. **B-C)** Female *Scn1a*^*+/-*^ mice are more susceptible to dying at both juvenile (<P30) and adult ages with mortality rate (slope) consistently increased throughout life compared to male mice. Mortality at P30: Male = 17.3%, Female = 30.8%; N = 120-125 *Scn1a*^*+/-*^, 56-70 *Scn1a*^*+/+*^. Log-rank test, Chi-square test, Simple linear regression or unpaired t-test, *p < 0.05, ****p < 0.0001, or p-value indicated; Relative risk (95% CI) = 1.19 (1.03 to 1.39).

Mice surviving to adulthood (>P90) were subjected to video-EEG for seizure monitoring. Converse to the mortality findings, more adult male *Scn1a*^*+/-*^ mice exhibited GTC seizures than female mice (Fig 2A). Additional sex divergences were seen in seizure characteristics, including seizure frequency, total ictal time and average seizure duration (Fig 2B-D). These were all increased in male *Scn1a*^*+/-*^ mice relative to female mice, suggesting higher seizure severity.

**Figure 2:**
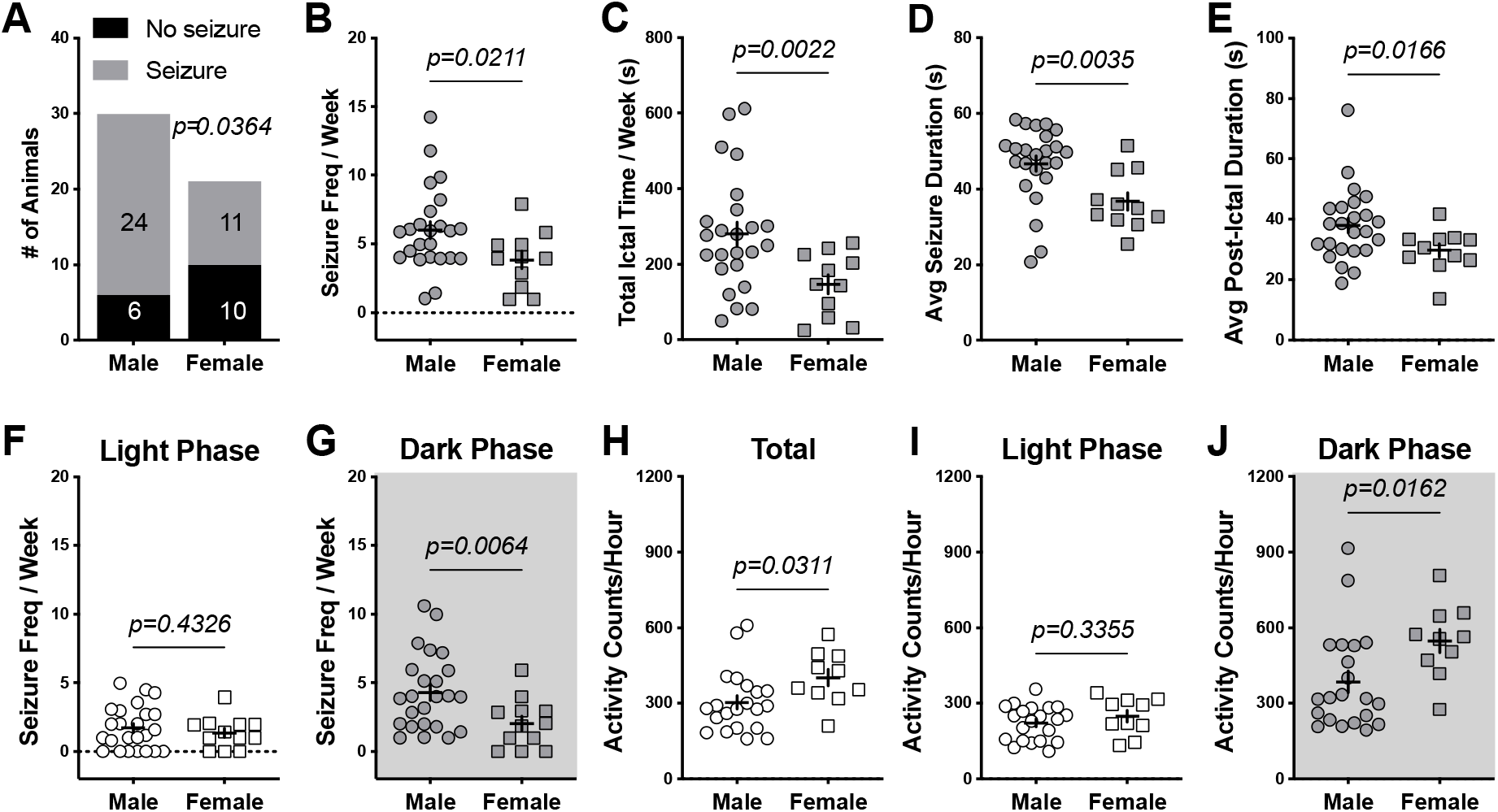
Adult male *Scn1a*^*+/-*^ mice exhibit more and severer convulsive seizures but hypoactive than female mice. **A)** Adult (P90) male *Scn1a*^*+/-*^ mice have more seizures than their female counterparts. **B-E)** In animals with seizures, seizure frequency, total ictal time, average (avg) seizure duration and avg post-ictal duration is greater/longer in male than female *Scn1a*^*+/-*^ mice. **F-J)** Seizing male *Scn1a*^*+/-*^ mice exhibit more seizures at night and are hypoactive compared to female mice. Seizure occurrence: Male = 80.0%, Female = 52.4%. N = 30 Male, 21 Female. Chi-square test or unpaired t-test with Welch’s correction, p-value indicated; Relative risk (95% CI) = 0.42 (0.18 to 0.95).

Therefore, despite the constant mortality increase in female *Scn1a*^*+/-*^ mice even at adulthood, morbidity appeared to be more severe in male *Scn1a*^*+/-*^ mice. Perhaps recovery from seizures was longer in female *Scn1a*^*+/-*^ mice; however, duration of post-ictal period was also increased in male *Scn1a*^*+/-*^ mice (Fig 2E). Given the heightened rate of seizures and SUDEP at night (13,14), we further examined seizure frequency in each diurnal phase. As shown in Figure 2F-G, no sex difference in seizure frequency was apparent during the day; but, male *Scn1a*^*+/-*^ mice exhibited more seizures at night than female mice. We then examined home-cage locomotor activity as an approximate indicator of sleep given previous studies of sleep deficits in *Scn1a* mutant mice (7,15), and the inverse relationship of sleep with seizure incidence and SUDEP (7,13). Accordingly, we found that seizing female *Scn1a*^*+/-*^ mice exhibited hyperactivity compared to seizing male mice overall and at night (Fig 2H-J). This increased activity suggests increased wakefulness and sleep disruption, which may contribute to a higher risk of mortality in female *Scn1a*^*+/-*^ mice.

## Discussion

Sex differences in phenotype presentation have implications in therapeutic assessment in preclinical and clinical studies. We sought to determine if sex differences in disease phenotype, partly previously identified in juvenile *Scn1a*^*+/-*^ mice, are observed in adult animals. Our results in overall mortality for juvenile (<P30) and adult (P30-90) *Scn1a*^*+/-*^ mice are in agreement with previous findings (10,12). The observed increased mortality in female *Scn1a*^*+/-*^ animals regardless of age is however a novel finding. Our findings are consistent with clinical reports of female DS patients succumbing to seizure-related deaths independent of age (3,5), further highlighting the clinical relevance of these sex difference findings and the validity of this mouse model.

Given the increased incidence of SUDEP in female *Scn1a*^*+/-*^ mice, it is not surprising that seizure susceptibility is also increased in young animals as previously observed during the worsening (P19-30) and/or stabilization (P36-49) stages (10,12). However, contrary to these findings, we observed increased seizure susceptibility and severity (seizure number and duration) in adult male *Scn1a*^*+/-*^ animals. This suggests that the female *Scn1a*^*+/-*^ mice that survive the juvenile mortality are less susceptible to seizures. It has been previously stipulated that intrinsic sex differences in inhibition may account for the observed distinct phenotypic presentation (10). But again, in contrast to our findings, these differences point to female mice having more seizures. For example, adult female DS mice with dysfunction in GABAergic signaling via a GABA_A_ receptor mutation have more epileptic spike-wave discharges (SWDs) (16). This raises the possibility that abnormal SWDs, typically seen in absence epilepsy, are more common in females, and their occurrence, compounded with GTCs, increases mortality. Abnormal SWDs have not been thoroughly investigated in *Scn1a*^*+/-*^ mice, and are not present in our studies with adult mice. This is consistent with the incidence of absence epilepsy restricted to the worsening stage in patients (9). Still yet, such possibilities of intrinsic sex differences need to be empirically tested.

The observation of increased mortality with decreased seizure incidence and morbidity in adult female *Scn1a*^*+/-*^ mice is an interesting conundrum. The shorter post-ictal recovery observed in female mice contrasts with studies implicating longer recovery from seizures as a SUDEP risk (17,18). This however does not eliminate post-ictal recovery as a SUDEP contributor given that respiratory and cardiovascular dysfunction are not specific to the post-ictal period. Kalume *et. al*. for example showed that ictal bradycardia is common in juvenile female *Scn1a* mutant mice after hypothermia-induced seizures (7). The brainstem may account for these changes in autonomic function as previously reported. Specifically, in *Scn1a* mutant mice, viral-mediated *Scn1a* re-expression in the brainstem and cerebellum increased hypothermia-induced seizure threshold and survival (12,19).

Along these lines of the brainstem’s role in SUDEP, we explored the factor of sleep/activity in influencing sex-biased mortality. Studies with DS mice and patients have demonstrated that sleep disturbances increases seizure incidence and SUDEP risk (7,9,15,16). Of note, *Scn1a* mutant mice exhibit increased wakefulness in the dark phase (15), which we also observed in female *Scn1a*^*+/-*^ mice compared to their male counterparts in the form of hyperactivity. This sleep/activity dysfunction could therefore impact seizure-related mortality that is increased in female *Scn1a*^*+/-*^ mice. However, hyperactivity does not translate to wakefulness. Thus, more detailed sleep studies are needed to elucidate the relationship between sleep, SUDEP and sex in these mice.

Overall, we found a significant sex and age divergence of mortality and epilepsy profiles in *Scn1a*^*+/-*^ mice. Further studies are still needed to understand the mechanisms underlying these differences. Nonetheless, our findings underscore the potential confounders and biases during screens for anti-epileptic interventions using this model and highlight the need to monitor and consider these factors in studying DS and *SCN1A*-related disorders.

## Acknowledgements

We thank the IDDRC Animal Behavior and Physiology Core at Boston Children’s Hospital, funded by NIH/NICHD P50 HD105351.

## Author Contributions

Conceptualization & Methodology: S.A.D.V., M.Q.H. and A.R.; Formal analysis and Investigation: S.A.D.V., R.L., G.M., A.L., Y.C., B.H., E.L., S.C.D, and M.Q.H.; Data curation: S.A.D.V.; Resources: A.R.; Writing: All authors; Visualization: S.A.D.V., M.Q.H. and A.R; Supervision: A.R.

